# The Multispecies Coalescent Model Outperforms Concatenation across Diverse Phylogenomic Data Sets

**DOI:** 10.1101/860809

**Authors:** Xiaodong Jian, Scott V. Edwards, Liang Liu

**Author notes:** Corresponding authors Scott Edwards, Department of Organismic and Evolutionary Biology Harvard, Cambridge, MA, Liang Liu, Department of Statistics Institute of Bioinformatics University of Georgia.

## Abstract

A statistical framework of model comparison and model validation is essential to resolving the debates over concatenation and coalescent models in phylogenomic data analysis. A set of statistical tests are here applied and developed to evaluate and compare the adequacy of substitution, concatenation, and multispecies coalescent (MSC) models across 47 phylogenomic data sets collected across tree of life. Tests for substitution models and the concatenation assumption of topologically concordant gene trees suggest that a poor fit of substitution models (44% of loci rejecting the substitution model) and concatenation models (38% of loci rejecting the hypothesis of topologically congruent gene trees) is widespread. Logistic regression shows that the proportions of GC content and informative sites are both negatively correlated with the fit of substitution models across loci. Moreover, a substantial violation of the concatenation assumption of congruent gene trees is consistently observed across 6 major groups (birds, mammals, fish, insects, reptiles, and others, including other invertebrates). In contrast, among those loci adequately described by a given substitution model, the proportion of loci rejecting the MSC model is 11%, significantly lower than those rejecting the substitution and concatenation models, and Bayesian model comparison strongly favors the MSC over concatenation across all data sets. Species tree inference suggests that loci rejecting the MSC have little effect on species tree estimation. Due to computational constraints, the Bayesian model validation and comparison analyses were conducted on the reduced data sets. A complete analysis of phylogenomic data requires the development of efficient algorithms for phylogenetic inference. Nevertheless, the concatenation assumption of congruent gene trees rarely holds for phylogenomic data with more than 10 loci. Thus, for large phylogenomic data sets, model comparison analyses are expected to consistently and more strongly favor the coalescent model over the concatenation model. Our analysis reveals the value of model validation and comparison in phylogenomic data analysis, as well as the need for further improvements of multilocus models and computational tools for phylogenetic inference.

---

Due to the increasing growth in dimensionality and complexity of multilocus sequence data, accurate phylogenetic inference for understanding the evolutionary history of species faces substantial computational and modeling challenges (Rannala and Yang 2008; Liu *et al*. 2015; Edwards *et al*. 2016). With the assumption of topologically congruent (TC) genealogies across loci, a phylogenetic tree estimated from concatenated sequences is often used as the estimate of the species tree, despite the fact that the concatenation model oversimplifies the complexity inherent in the diversification of species by ignoring many biological phenomena, such as deep coalescence, hybridization, recombination, and gene duplication and loss, that are commonly observed during the history of species (Maddison 1997; Rannala and Yang 2003; Bravo *et al*. 2019). Since the advent of multilocus sequence data, there has been ongoing effort toward building stochastic models for handling gene tree variation. Given that incomplete lineage sorting (ILS) is likely the most common biological source of gene tree variation (Edwards 2009), some of the earliest efforts included parsimony and Bayesian multispecies coalescent models based on a coalescence process running along the lineages of the species tree (Page 1998; Liu and Pearl 2007; Liu 2008; Heled and Drummond 2010). Subsequently, multispecies coalescent (MSC) model has been updated by adding biological parameters, such as gene flow, rate variation among lineages, recombination and hybridization (Kubatko 2009; Hey 2010; Wang *et al*. 2014). In other developments, methods have been proposed to ask whether a tree is the best model for a given data set, or whether reticulations in the form of gene flow or lineage merging are more appropriate (see below) (Moret *et al*. 2004; Jackson *et al*. 2017; Burbrink and Gehara 2018). Meanwhile, skeptics of the MSC and advocates for simpler models, particularly concatenation models, have raised critical questions about appropriateness of the MSC and argued that observed gene tree variation is often not caused by ILS, but instead by gene tree estimation errors (Springer and Gatesy 2016; Scornavacca and Galtier 2017; Richards *et al*. 2018). Given the inconsistency of concatenation methods under some regions of tree space in which coalescent methods of tree building are still consistent (Liu *et al*. 2010), to many researchers it is the convincing evidence to choose coalescent over concatenation methods for species tree inference. However, since the mathematical proofs or simulations for inconsistency of concatenation methods assume that the MSC model is true (Kubatko and Degnan 2007; Roch and Steel 2015), it is important to show empirical evidence for validating and comparing the concatenation and MSC models.. Resolution of debates over concatenation and coalescent models requires a statistical framework of model comparison and model validation on a variety of empirical and simulated data.

## Model fit and model adequacy in phylogenomics

Discussions about the necessity or adequacy of MSC models versus concatenation have taken three principle forms. One form involves questioning the phylogenetic signal in data sets designed for application of the MSC. For example, several authors have suggested that gene tree error (whether deriving from alignment artifacts, low signal, or gene tree estimation error), rather than ILS, is responsible for most if not all observed gene tree variation (Gatesy and Springer 2014; Arcila *et al*. 2017). If demonstrable gene tree variation can be ruled out for a given data set, this logic goes, then concatenation is a reasonable fallback model for analysis. This logic reasonably implies that the simpler model inherent in concatenation is favored, especially when gene tree variation can be shown to be low or negligible. The problem with this logic, however, is that gene tree variation is only one motivation for MSC models. The other, more fundamental, motivation for MSC models is the conditional independence of loci in the genome, wherein recombination and random drift render the topologies, but more often the branch lengths of different loci independent of one another, conditional on the species tree, which necessarily influences the shape of all gene trees in the genome (Edwards *et al*. 2016). This point leads to the second common argument against MSC models: that violation of MSC model assumptions, such as evidence for recombination within loci or lack of recombination between loci or violations of neutrality, render MSC models poor descriptors of actual data sets, again recommending concatenation or other approaches as more robust alternatives (Gatesy and Springer 2014; Scornavacca and Galtier 2017). Again, however, demonstration of violation of a model’s assumptions does not necessarily imply that that model is not a reasonable, or even the best available, description of the data. Indeed, we know of no violation of assumptions of an MSC model that is not also a violation of the concatenation model, especially since the concatenation model is best described as a special case of the MSC model, wherein all gene trees are topologically identical (Liu *et al*. 2015).

A third approach to deciding whether MSC, concatenation or other models might best apply to a given data set is testing for model fit and model adequacy (Brown and Thomson 2018). Only a few papers have explicitly tested the fit and adequacy of the MSC, and, crucially, in doing so, have usually neglected to compare the fit of the major alternative model, concatenation, to that of the MSC. Reid et al. (Reid *et al*. 2014) applied posterior predictive simulation (PPS), a Bayesian modeling approach, to a series of moderately sized data sets and concluded that a “poor fit to the multispecies coalescent is widely detectable in empirical data.” Although this study was a major advance in our understanding of the MSC as applied to real data sets, we wonder whether the sweeping nature of this conclusion is reasonable and suggest that it may have been overly pessimistic. First, they definitively rejected the fit of the MSC at the level of gene trees for only 4 out of 25 data sets, and only 7 total loci (2.9%), hardly suggesting that poor fit at the level of the coalescent is “widespread”. Reid et al. (2014) also suggest that a large percentage of partitions or loci in data sets, sometimes as high as 50%, violate the MSC; but the largest data set in their analysis consisted of only 20 loci, with 15 out of 25 data sets consisting of less than 10 loci, thereby possibly exaggerating the extent of violations of the MSC. Many of these rejections were based not on a poor fit of coalescent assumptions but on deviations of the loci from assumed substitution models, which is hardly a direct rejection of the MSC itself. In several cases, Reid et al (Reid *et al*. 2014) were unable to distinguish whether the MSC was a poor fit due to analyses at the level of estimating gene trees and substitution models or at the level of the coalescent model itself. Additionally, their use of a χ^2^ test for comparing the observed site patterns with those expected from the assumed substitution models could not accommodate missing data, making this aspect of their model testing problematic and raising the possibility that different substitution models could improve model fit. Moreover, several of the data sets in their analysis can be reasonably thought of as phylogeographic data sets (e.g., Leache (2009) and Walstrom et al. (2012)), in which gene flow is likely or expected, rather than phylogenomic data sets, in which gene flow is less expected or unexpected. Arguably, none of the data sets they analyzed could be called robustly phylogenomic in the modern sense: only 5 of the data sets interrogated relationships above the level of genus and none of them examined relationships among families suborders or orders, where, again, gene flow among lineages is less expected. It is therefore perhaps not surprising that such data sets do not conform well to the MSC. Jackson et al. (2017) recently applied a novel model fitting algorithm, Phrapl, to a series of multilocus phylogeographic data sets, concluding that a pure isolation model, such as the MSC, is rejected in favor of models including gene flow and other reticulate events. Here, again, however, the data sets analyzed are explicitly termed phylogeographic, and, although the MSC was often applied to these data sets in the original papers, it is unsurprising that this model is a poor fit compared to models that include gene flow. Like all models, the MSC has a particular domain of application, one that we suggest is even wider than that in which concatenation is appropriate, but not so wide as to be applicable to data sets that demonstrably include gene flow or hybridization.

Most arguments in favor of concatenation, such as those summarized above, assert the superiority of concatenation by noting widespread perceived violations of the MSC or lack of demonstrable gene tree variation, and there have only been a few scattered examples of statistical comparisons of model fit and model adequacy between the MSC and concatenation. To our knowledge, only Liu and Pearl (2007) and Edwards et al. (2007) have explicitly compared the fit of MSC and concatenation models to the same data set, and asked which is a better fit (model comparison) and whether either model can account for the details of the multilocus data (model adequacy). Tests of model fit and model adequacy have been conspicuously absent from discussions about the relative merits of the MSC versus concatenation, although they have begun to appear in discussions of the relative merits of simple and more elaborate MSC models (Wen *et al*. 2016; Jackson *et al*. 2017). Testing whether the MSC or concatenation can adequately describe multilocus data sets, and which model can describe those data better, will go a long way towards addressing concerns about the MSC and towards delimiting its appropriate domain of application.

In this paper, we will address several questions regarding the MSC and its application to empirical data sets: (1) is gene tree variation primarily caused by estimation error? (2) how well do the assumed substitution models fit multilocus sequence data and how does this fit drive the overall fit of the MSC? (3) how well does the multispecies coalescent model, divorced from shortcomings of the substitution model, fit empirical data? and (4) which model fits empirical multilocus data sets better, concatenation or the MSC? Question 1 can be addressed by a likelihood ratio test (LRT), where the null hypothesis is that all loci have the same tree topology (but possibly different branch lengths) versus the alternative hypothesis that allows gene tree variation across loci (testing TC gene trees; Figure 1). We addressed question 2 with the same χ^2^ test used by Reid et al. (Reid *et al*. 2014), comparing the observed and expected frequencies of site patterns, but this test cannot be applied to sequences with missing characters. We therefore modified this τ^2^ test to handle missing characters (goodness of fit of substitution models; Figure 1). To evaluate the effect of substitution models on the fit of the MSC, model validation analysis was conducted with different substitution models for the same data sets. Question 3, the adequacy of the MSC, can be addressed using posterior predictive simulation (PPS) in a Bayesian framework (Reid *et al*. 2014) implemented in a Bayesian phylogenetic program BEAST (Suchard *et al*. 2018). Unlike previous studies utilizing sequence data simulated under the MSC model to validate coalescent methods (Kubatko and Degnan 2007), PPS directly evaluates the fit of the MSC model by comparing the posterior gene trees and the gene trees simulated from the MSC (Reid *et al*. 2014). However, the posterior gene trees generated with a coalescent prior are biased towards the MSC, especially when the alignments lack phylogenetic signal and Bayesian inference of gene trees is primarily driven by the coalescent prior. Validation analysis should instead compare the simulated gene trees with empirical gene trees estimated from multilocus sequence data without any influence of the MSC (i.e., the posterior gene trees generated independently across loci). Additionally, to reduce the influence of substitution models on the fit of the MSC, we perform the model validation only for loci that fit the substitution model (Bayesian model validation; Figure 1). Question 4, model comparison between the concatenation and MSC models, can again be addressed in a Bayesian framework, using Bayes factors (Bayesian model comparison; Figure 1) or other posterior predictive approaches (Lewis *et al*. 2014). By addressing these questions, we aim to directly compare competing models, especially as they apply to phylogenomic data sets where concatenation might plausibly be applied.

**Figure 1:**
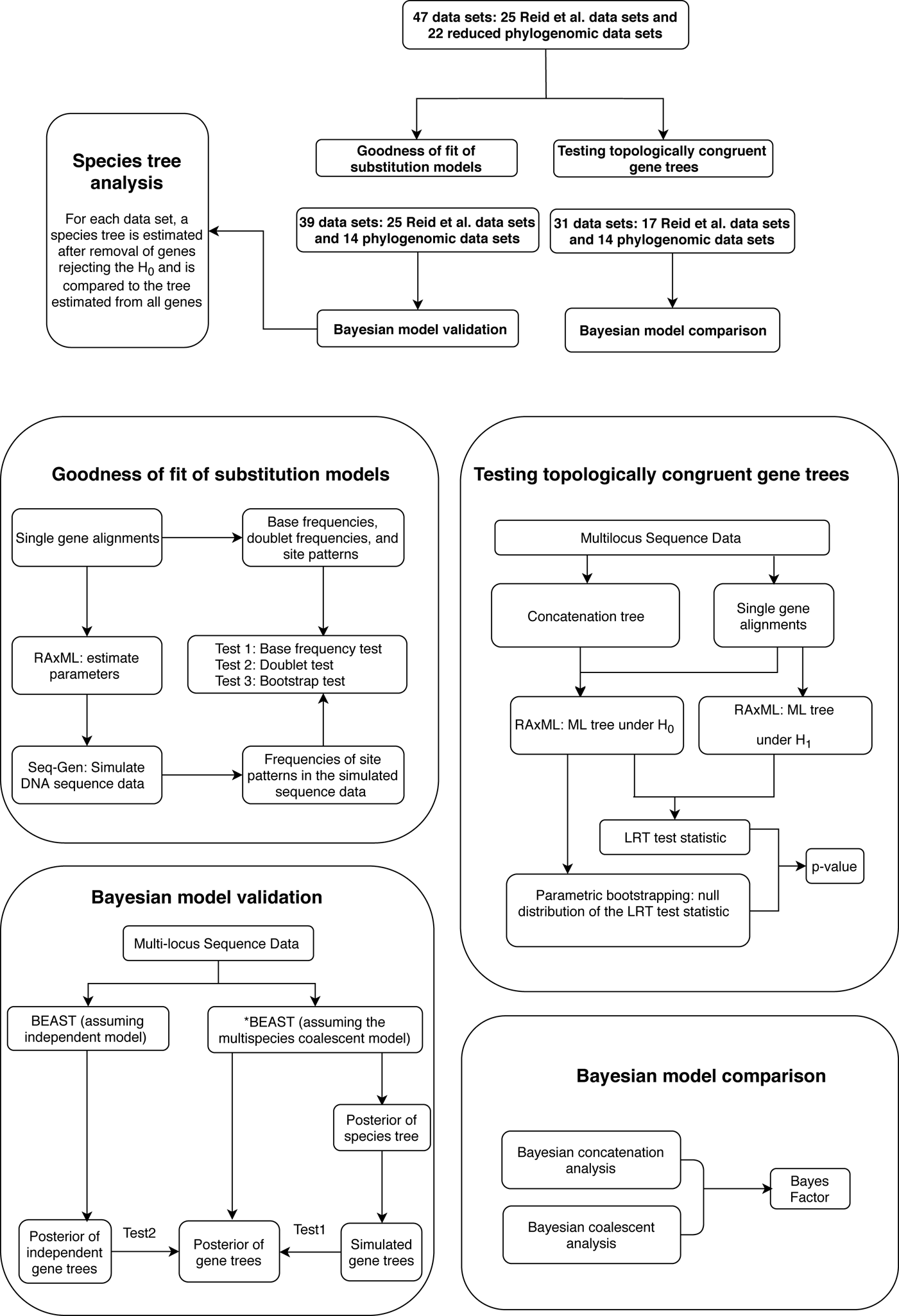
The flowchart of phylogenetic analyses in this study. The alignments of 47 data sets were used to test the hypothesis of topologically congruent (TC) gene trees and goodness of fit of substitution models. The model validation and comparison analyses were only performed for the loci that fit the GTRGAMMA model, reducing the number of input data for model validation to 39 (25 Reid et al. data sets and 14 phylogenomic data sets). Because the model comparison analysis implemented in BEAST does not allow missing taxa in any locus, 8 Reid et al. data sets were removed, further reducing the number of input data for the model comparison analysis to 31 (17 Reid et al. data sets and 14 phylogenomic data sets).

## MATERIALS AND METHODS

### Phylogenomic data sets

This study consists of 47 empirical data sets, including 25 data sets from Reid et al (Reid *et al*. 2014) and 22 phylogenomic data sets across the tree of life. We chose phylogenomic data sets primarily based on their having been sampled from multiple species, usually more than 10 at the level of family or above, for their coverage of at least 50 loci and for their availability in already aligned nexus or phylip format on an easy accessible open access data base. We also eased our search by focusing primarily although not exclusively on data sets from the journal Molecular Phylogenetics and Evolution. The 22 phylogenomic data sets were downloaded from the data links available in the original papers (Table 1). The genetic markers of 22 phylogenomic data sets are highly diversified, including CDS matrices, exons, and UCEs (Table 1). There are 12 to 207 species in the 22 additional data sets, and the number of loci ranges from 110 to 30,636 (Table 1). Due to computational limits of Bayesian phylogenetic analyses, each of the 22 phylogenomic data sets was reduced to only include the alignments of the 10 most fully populated species across loci, and loci with missing sequences were removed from further analysis. Removing species from alignments will, if anything, increase gene tree similarity across loci as compared with data sets with the full complement of species, and therefore increase the fit of the concatenation model to the data. After data reduction, the data sets contained alignments of 36 (Aitken *et al*. 2017) to 4709 (Wu *et al*. 2018) loci, each with 10 species (Table 1).

**TABLE 1:** Summary of 22 phylogenomic data sets analyzed here. In the columns OTUs and Loci, the bottom number is the count of taxa (or loci) in the original data set; the top number is the count of taxa (or loci) after data reduction.<colcnt=2>

| Data set | OTUs | Loci | Markers |
| --- | --- | --- | --- |
| Ant (BLAIMER *et al.* 2018b) | 10/153 | 1509/1763 | UCE |
| Bee (SANN *et al.* 2018) | 10/95 | 193/195 | CDS |
| Bird (PRUM *et al.* 2015) | 10/198 | 259/259 | CDS |
| Lizard (BLOM *et al.* 2017) | 10/29 | 1831/1852 | Exon |
| Brittlestar (O'HARA *et al.* 2019) | 10/46 | 407/416 | CDS |
| Butterfly (ESPELAND *et al.* 2018) | 10/207 | 325/352 | Exon |
| CarpenterBee (BLAIMER *et al.* 2018a) | 10/179 | 597/753 | UCE |
| Cichlid (MCGEE *et al.* 2016) | 10/50 | 924/1043 | UCE |
| Clupeocephalan (STRAUBE *et al.* 2018) | 10/52 | 48/829 | Exon |
| Cracids (HOSNER *et al.* 2016) | 10/23 | 430/430 | UCE |
| Darter (MACGUIGAN AND NEAR 2018) | 10/112 | 363/30636 | Exon |
| Diplostomoidea (LOCKE *et al.* 2018) | 10/12 | 324/517 | UCE |
| Gallopheasant (MEIKLEJOHN *et al.* 2016) | 10/18 | 1479/1479 | UCE |
| Hemimastigophora (LAX *et al.* 2018) | 10/61 | 239/351 | CDS |
| Lepanthe (BOGARÍN *et al.* 2018) | 10/33 | 334/433 | CDS |
| Mammal2018 (WU *et al.* 2018) | 10/90 | 4709/5162 | CDS |
| Mammal2017 (SCORNAVACCA AND GALTIER 2017) | 10/97 | 108/110 | CDS |
| Metazoan2015 (WHELAN *et al.* 2015) | 10/76 | 36/115 | CDS |
| Shrew (GIARLA AND ESSELSTYN 2015) | 10/19 | 966/1112 | UCE |
| Squirrel (COOK *et al.* 2018) | 10/74 | 3209/3951 | UCE |
| Weevil (AITKEN *et al.* 2017) | 10/67 | 318/521 | CDS |
| Fish (CUI *et al.* 2013) | 10/27 | 1183/1183 | CDS |

The sequence divergence (i.e., the average pairwise p-distance) of the 22 reduced phylogenomic data sets was significantly higher (p-value < 0.01) than that of 25 Reid et al. data sets (Figure A1), indicating that these new data sets are more classically ‘phylogenomic’ than those of Reid et al. (2014). Indeed, several of the data sets we analyze are at very high taxonomic levels, such as those of metazoans (Whelan *et al*. 2015; Simion *et al*. 2017) or mammals (Scornavacca and Galtier 2017; Wu *et al*. 2018). Some of these data sets, such as those from metazoans, are at such high taxonomic levels, that MSC models have to our knowledge never been applied, perhaps in the mistaken idea that ILS is extremely unlikely to occur among such deep lineages. However, as pointed out by many authors, even if ILS is harder to detect among deep lineages, it is no less likely to occur among deeply diverging than recently diverging lineages, since it is the length of internodes, rather than their depth in time, that is most relevant to MSC processes.

### Goodness of fit of substitution models

Let *X = {x_1_,…, x_k_}* be the counts of *k* site patterns in a locus alignment. The random variables *X = {x_1_,…, x_k_}* have a multinomial distribution with 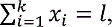 where *l* denotes the length of the alignment. Let *M* be the pre-selected substitution model. When there is no missing data, the τ^2^ goodness of fit test can evaluate adequacy of model *M* by comparing the observed and expected counts of site patterns (Reeves 1992; Goldman 1993; Jhwueng 2013). The test statistic is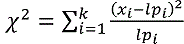, in which is the probability of site pattern *i*, which can be estimated under the null hypothesis using the maximum likelihood (ML) estimates of the tree topology, branch lengths, and the parameters in model *M*. Asymptotically, the test statistic has the τ^2^ distribution with (*k-*1) degrees of freedom. Since this τ^2^ test statistic is based on the frequencies of sites for fully populated characters, sites with missing characters (gaps, ambiguous nucleotides, and unidentified regions) are excluded from analysis, even though only a small portion of nucleotides are missing in each site. To incorporate partially missing sites, a modified τ^2^ test was developed to calculate the observed and expected counts of site patterns in the presence of missing data (Waddell 2005). We instead calculate the marginal proportion of a site with missing characters and then compare the marginal proportion of the site with its expectation. For example, there are two missing characters in a site {??AC} of 4 species (*S_1_, S_2_, S_3_, S_4_*), i.e., the nucleotides from species *S_1_* and *S_2_* are missing. The marginal proportion *}_AC_*, of {AC} in the alignment of *S_3_* and *S_4_* is given by *}_AC_,= x_AC_/z*, in which *x_AC_* is the count of {AC} and *z* is the number of sites without missing characters in the alignment of *S_3_* and *S_4_*. Since a small *z* indicates a large amount of missing data in the alignment, we ignore the sites for which *z*/*l* ≤ 0.8. The test statistic 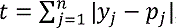 in which is the observed proportion of a site pattern with or without missing characters, and *p_j_* is the probability of the site pattern under the null hypothesis, and *n* is the number of site patterns for which *z*/*l* > 0.8. The null distribution of the test statistic is estimated by parametric bootstrap samples generated from the ML estimates of the tree topology, branch lengths, and the parameters in model *M*.

Marginal tests of base compositions between pairs of taxa (Tavare 1986) have been found to be more powerful than the τ^2^ goodness of fit test of the substitution models to phylogenetic data (Waddell *et al*. 2009). We therefore perform two marginal tests. The first test is to detect heterogeneity of base compositions across species, when the observed base frequencies of individual species deviate significantly from the overall average base frequencies across species. Let {x_A_, x_C_, x_G_, x_T_} be the observed frequencies of nucleotides A, C, G, and T of a species. The frequency of nucleotide *i (i = A, C, G, T)* is equal to *x_i_ = N_i_/N*, where *N_i_* is the number of nucleotide *i* and *N* is the total number of nucleotides excluding missing characters in the sequence. Under the null hypothesis of homogeneous substitution models, all species are expected to have the same base frequencies and thus the expected base frequencies under the null hypothesis are estimated by the overall average frequencies {p_A_, p_C_, p_G_, p_T_} of A, C, G, and T across species. The τ^2^ test of the observed frequencies {x_A_, x_C_, x_G_, x_T_} against the expected frequencies {p_A_, p_C_, p_G_, p_T_} is carried out for each species to detect those species whose base frequencies significantly deviate from the null hypothesis that all species have the same base frequencies. Similarly, the second marginal test is applied to the frequencies of double-nucleotides (doublets) between pairs of species to find pairs of species for which doublet frequencies are inconsistent with the pre-selected substitution model. The frequency of doublet *ij(i = A, C, G, T and j = A, C, G, T)* is equal to *x_ij_ = N_ij_/N*, is the number of doublet *ij* and *N* is the total number of doublets excluding those with missing characters.

The bootstrap and two marginal tests were conducted to evaluate goodness of fit of the GTRGAMMA model to 47 empirical data sets (Figure 1). We chose the GTRGAMMA model because the most parameter-rich model (GTRGAMMA in RAxML) is sufficient for reliable phylogenetic inference (Abadi et al. 2019) and a more complex model tends to be a better fit to phylogenomic data (Liu *et al*. 2017). For each locus, the ML estimates of the phylogenetic tree and other parameters were obtained by RAxML v8.2.3 (Stamatakis 2014) using the GTRGAMMA model. Then, 100,000 base pairs of sequence were simulated from the estimated phylogenetic tree using Seq-Gen v1.3.2x (Rambaut and Grassly 1997). The expected frequencies of site patterns were estimated by the corresponding frequencies in the simulated sequences. In addition, 100 bootstrap samples were generated by simulating DNA sequences from the concatenation tree for each locus. If taxa were missing from the loci, they were pruned from the concatenation tree and DNA sequences were simulated from the pruned concatenation tree. The test statistic was calculated for each bootstrap sample, and the observed test statistic t* was compared with the bootstrap test statistics *{t_1_,…,t_100_}* and p-value = (# of t_1_ > t*) / 100. For the marginal test of base frequencies, the sequence of a species was considered significant if its p-value was less than 0.05 divided by the number of species. A locus was significant if it had at least one significant species. Similarly, for the test of doublet frequencies between pairs of species, a pair of species was significant if its p-value was less than 0.05 divided by the number of pairs. A locus was significant if it has at least one significant pair. The bootstrap and two marginal tests were applied to each locus of the 47 data sets. A locus waas significant if any of above three tests was significant for the locus.

### Testing for topologically congruent gene trees

We ask the question whether a single gene tree topology can adequately explain a given multilocus data set, using a variant of the LRT. Similar types of LRTs were developed to test if alternative trees are congruent with the maximum likelihood (ML) tree for a single locus (Shimodaira and Hasegawa 1999; Shimodaira 2002). Mcvay and Carstens (2013) proposed a parametric bootstrap approach to assess the extent to which gene tree variation can be attributed to phylogenetic estimation error. Here, we develop a LRT to evaluate the concatenation assumption of congruent gene trees for multiple loci. Let D = (D, D_;5_,…, D) be the concatenated alignments of a multilocus sequences data set, in which D represents the alignments of locus *i* and *K* is the count of loci. The tree topology of locus *i* is denoted by 1.. Under the concatenation model, all loci are assumed to have the same tree topology, i..e, τ_1_ = … = τ_K_. We develop an LRT to evaluate the null hypothesis that all gene trees have the same topology, i.e., τ_1_ = … = τ_K_. versus the alternative hypothesis that not all gene trees are topologically identical. The test statistic is defined as t = log(l_1_) - log(l_0_), in which *l_0_* and *l_1_* are the likelihoods of the null and alternative hypotheses. Under the null hypothesis, the ML tree is built from the concatenated alignments across loci using RAxML v8.2.3 (Stamatakis 2014) with the GTRGAMMA model (Tavare 1986; Yang 1994). Using the ML tree instead of the true concatenation tree for the null hypothesis may lead to a biased test. However, since the concatenated sequences of phylogenomic data consist of millions of base pairs, the ML tree is very similar, if not identical, to the true concatenation tree. hus, the bias induced by the topologcial difference between the ML and true concatenation tree is negaligible for phylogenomic data. Let N\ be the log-likelihood of locus *i* by refitting branch lengths and substitution model parameters to the concatenation tree with missing taxa being removed. The log-likelihood under the null hypothesis is equal to the sum of the log-likelihoods of individual loci, i.e., 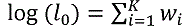. Refitting branch lengths and model parameters on a fixed tree topology was performed in RAxML using the command line “*raxml-HPC-AVX –s datafile –m GTRGAMMA –n outputfile –f e –t fixtree*”. To find the log-likelihood of the alternative hypothesis, maximum likelihood trees were independently built for individual loci using RAxML. Since model parameters include tree topologies, the τ^2^ distribution is not a good approximation to the null distribution of the test statistic (Jhwueng *et al*. 2014). Therefore, the null distribution of the test statistic was approximated by a parametric bootstrap. Bootstrap samples were generated under the null hypothesis by simulating DNA sequences from the concatenation tree pruned for available species at each locus. Since the original alignments include missing characters (gaps, ambiguous nucleotides, and unidentified regions), bootstrap samples should involve a similar pattern of missing characters. Thus, corresponding nucleotides in bootstrap samples were replaced by missing characters. Let \: be the value of the test statistic *t* for the bootstrap sample *i* = 1, …, *B*. The loglikelihoods of the null and alternative hypothesis in the test statistic *t* were generated using RAxML. The values *{t_i,_ i = 1, …, B}* of the test statistic for *B* bootstrap samples were used to approximate the null distribution of the test statistic *t*. The p-value was estimated by the proportion of *{t_i,_ i = 1, …, B}* that were greater than or equal to the test statistic *t* calculated from real data. Rejection of the null hypothesis indicates that some gene trees are incongruent with the concatenation tree. Then, the LRT was further applied to each locus to identify alignments that reject the null (concatenation) tree. Let τ be the concatenation tree and τ_i_ is the gene tree *i*. The null and alternative hypotheses are H_0_: τ_i_ = τ and H_1_: τ_i_ ǂ τ. The test statistic is t = log(l_1_) - log(l_0_), in which log (l_0_) and log (l_1_) are the log-likelihoods of locus *i* under the null and alternative hypotheses. The null distribution of the test statistic was approximated by the bootstrap samples generated under the null hypothesis and the p-value was equal to the proportion of bootstrap test statistics that are greater than or equal to the observed test statistic. A locus was significant if its p-value was less than or equal to 0.05 divided by the number of loci (Bonferroni correction for multiple comparisons).

### Bayesian validation of the multispecies coalescent model

Like Reid et al. (2014), the Bayesian MSC model is here validated by Bayesian predictive simulation. Reid et al. recommended comparing the posterior coalescent gene trees to the gene trees simulated from the posterior species trees. However, the posterior coalescent gene trees have been influenced by the multispecies coalescent prior, which is implemented in packages such as BEST (Liu 2008) or *BEAST (Heled and Drummond 2010). The posterior gene trees with an MSC prior are biased towards the MSC model to some extent, compared to gene trees generated without an MSC prior, making this test conservative (i.e., in favor of the MSC). Thus, here, simulated gene trees are compared to posterior gene trees generated without the MSC prior. We validate the MSC model using two tests. The first test, as in Reid et al. (2014), involves two comparisons - comparing the multispecies coalescent likelihoods of the simulated and posterior coalescent gene trees and comparing the number of deep coalescences of the simulated and posterior coalescent gene trees. The first test is rejected if either or both of two comparisons reject the MSC model. In the second test, not used by Reid et al. (2014), the posterior gene trees are first estimated independently across loci in BEAST by unlinking the substitution models, clock models, and trees. If the 99.9% posterior credible regions of the log-likelihoods of the independent model and the MSC model are not overlapped, we conclude that there is a significant difference between the posterior independent and coalescent gene trees. We infer a poor fit of the MSC model to the data if either or both of the two tests are rejected. Both tests can also be applied to each locus in the data to identify gene trees that significantly deviate from the MSC model.

To alleviate the impact of poorly fitting substitution models on the fit of the MSC, the model validation and comparison analyses were only performed on the loci that fit the assumed substitution model (GTRGAMMA). Since the number of loci in 25 Reid et al. data sets is insufficient for filtering out the unfit loci, selection of loci that fit the substitution model was only performed on 22 reduced phylogenomic data sets. After locus selection, 9 of 22 data sets had ≤ 3 loci that fit the substitution model and thus they were removed from further analysis. We then randomly selected 50 loci from the remaining 14 phylogenomic data sets, resulting in a total of 39 data sets (25 Reid et al. data sets + 14 phylogenomic data sets with 50 loci) for the model validation analysis (Figure 1). The xml input files of the 25 data sets from Reid et al. were available in the data package provided by the authors. Reid et al. (2014)’s *BEAST analyses of those data sets assumed HKY and TN93 (+GAMMA) substitution models for individual loci and a multispecies coalescent prior for gene trees. To evaluate the effect of substitution models on the fit of the MSC model, Bayesian model validation of 25 Reid et al. data sets was reconducted with the GTRGAMMA model. For 14 phylogenomic data sets, to reduce the effect of substitution models on the overall fit of the MSC, the validation analysis was conducted only for loci for which the GTRGAMMA model was a good fit. Consequently, rejections of the MSC imply a poor fit of coalescent assumptions rather than a poor fit of the assumed substitution model. The xml input files of 14 data sets were generated using BEAUti v1.8.4, assuming the GTRGAMMA substitution model for all loci.

Two independent runs were carried out for each analysis and convergence was checked by comparing the outputs from two runs. The first 50% of MCMC samples were discarded as burn-in. Then, 1000 samples were selected from the remaining MCMC samples and used as input for Bayesian model validation. The first test of Bayesian model validation was conducted using the R package starbeastPPS (2014). To perform the second test, 39 input data sets were re-analyzed by unlinking the substitution models, clock models, and trees across loci in BEAST, which produced posterior gene trees independently across loci without the MSC prior. The 99.9% credible region of the difference between the log-likelihoods of the independent and coalescent models was calculated in R. The two tests were also applied to each locus to identify gene trees that significantly deviate from the MSC model.

To evaluate the effect of loci rejecting the MSC on species tree estimation, species trees were built from all loci and only loci that did not reject the MSC for each of the 14 phylogenomic data sets using NJst (Liu and Yu 2011) implemented in an R package Phybase (Liu and Yu 2010). This analysis was not performed for 25 Reid et al. data sets due to their small numbers of loci for species tree estimation. We calculated the Robinson-Foulds (RF) distance (Robinson and Foulds 1981) of two species trees reconstructed for each data set using the function dist.topo in an R package APE (Paradis and Schliep 2019). For the species trees with a positive distance, to evaluate statistical significance of the difference between two species trees, bootstrap support values of the incongruent branches were calculated using bootstrap gene trees estimated from alignments. Specifically, bootstrap gene trees were built for each locus and then used as input data to calculate bootstrap NJst trees.

The bootstrap support value of a branch was equal to the count of bootstrap NJst trees supporting the clade indicated by the branch. Romiguier et al. (2013) suggested that high GC content may cause problems for phylogenetic inference under the MSC model, and that selecting AT-rich loci can improve the resolution of estimated phylogenies. To investigate the association between GC content and poor fit of the MSC, we calculated GC content for loci rejecting and accepting the MSC, respectively. A two-sample t-test was used to find significant difference in GC content between loci rejecting the MSC and those accepting the MSC.

### Bayesian model comparison for MSC versus concatenation

Bayesian model comparison for concatenation versus the MSC was evaluated using Bayes factors (BF) (Kass and Raftery 1995), the ratio of marginal likelihoods BF = P(D|M_1_)/P(D|M_2_), in which *D* denotes data and M_1_ and M_2_ are two competing models. Here, *M_1_* is the MSC model and *M_2_* is the concatenation model. The Bayesian concatenation analyses assumed a partition model by unlinking substitution models and clock models across loci. The marginal log-likelihoods of the Bayesian multispecies coalescent and concatenation models were estimated using path sampling / stepping-stone sampling with 100 path steps implemented in BEAST. A value of log(BF) > 10 indicates that the Bayesian multispecies coalescent model *M_1_* is strongly favored by the data versus the Bayesian concatenation model *M_2_*.). The model comparison analysis was applied to the (39) data sets used in the Bayesian model validation analysis. Since the model comparison analysis implemented in BEAST does not allow missing taxa in any locus, 8 data sets with missing data were removed, further reducing the number of input data for the model comparison analysis to 31 (Figure 1).

To demonstrate that model assumptions may influence species tree inference, we reconstructed species trees for 5 phylogenomic data sets using concatenation and a coalescent method (NJst). In addition, we subsampled 25%, 50%, and 75% of loci from the original phylogenomic data, and compared the species trees built from the subsamples with those for the full phylogenomic data. The concatenation trees were estimated by RAxML with the GTRCAT model. The NJst trees were built using the function sptree.njst() in an R package Phybase. We calculated the bootstrap support values for each estimated species tree. Two estimated species trees are said to be significantly incongruent if two trees have a conflict branch with a bootstrap support value > 70. The subsampling analysis was repeated 10 times and we reported the proportion of subsamples (out of 10) for which the estimated species tree was significantly incongruent with the species tree built from the full phylogenomic data.

## RESULTS

### Goodness of fit of substitution models

There was a total of 20,032 loci (CDS, UCEs or exons) throughout the 47 empirical data sets; 241 loci from the 25 Reid et al. data sets and 19,791 loci from the 22 phylogenomic data sets. The marginal test of base frequencies identified a total of 1,362 (7%) loci/alignments (or 9% loci per data set in Figure 2a) for which at least one sequence significantly deviates from the average base frequencies expected from the GTRGAMMA model. The doublet test indicated that 6,990 (35%) loci/alignments (or 51% loci per data set in Figure 2a) have at least a pair of species whose doublet patterns are significantly different from the patterns expected from the GTRGAMMA model. The marginal test favors complex substitution models, because a complex model has a higher likelihood than a simple model, indicating that the expected frequencies under the complex model are more consistent with the observed frequencies of site patterns in the alignments. Thus, rejection of the GTRGAMMA model suggests that a more complex substitution model should be fit to the data. In the bootstrap test for site patterns, the GTRGAMMA model fails to fit the alignments of 5,718 (24%) loci across the 47 data sets (or 34% of loci per data set in Figure 2a). The doublet test appears to be more likely than the bootstrap test for site patterns to reject the GTRGAMMA model (Figure 2a), indicating the necessity of marginal tests for goodness of fit of substitution models. Because the doublet test is more likely to reject the GTRGAMMA model than the other two tests, the intersection test (the intersection of three tests, i.e., at least one of two marginal tests and the bootstrap test reject the GTRGAMMA model) appears to be primarily driven by the doublet test (Figure 2a). Overall, nearly half of the alignments (8,775 loci or 44%) were found significant (p-value < 0.05) in the intersection test, and the proportion of significant loci identified by the intersection test ranges from 6% to 100% across the 47 data sets (Figure 2a). A two-sample t-test finds no significant difference in the proportion of loci rejecting the GTRGAMMA model between the 25 Reid et al. data sets and the 22 phylogenomic data sets (Figure 2b).

**Figure 2:**
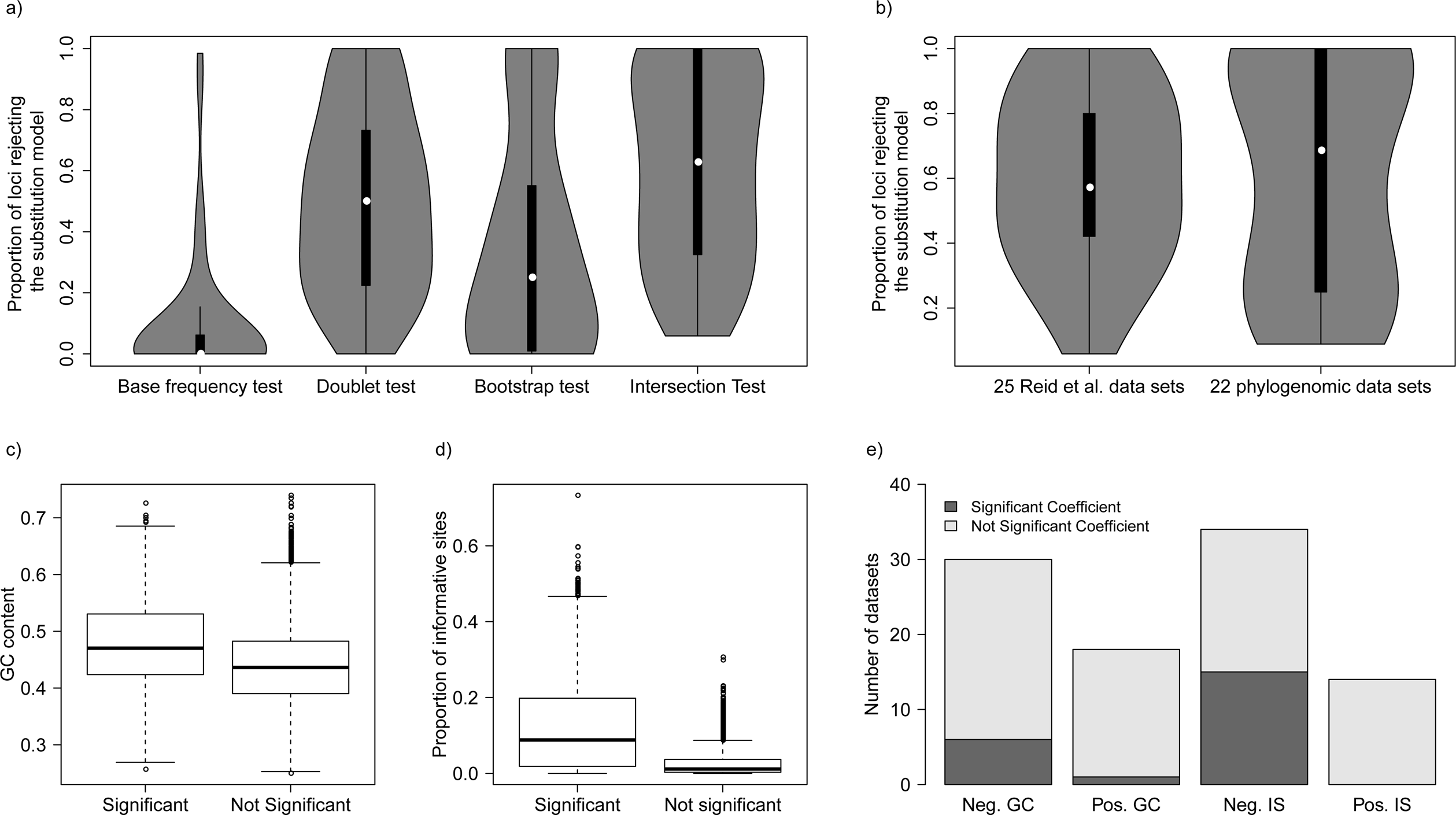
Goodness of fit of substitution models for 47 data sets. a) The violin boxplot of the proportion of loci rejecting the GTRGAMMA model in 47 data sets by the base frequency test, doublet test, bootstrap test, and the intersection of three tests (i.e., the intersection test), respectively. b) The violin boxplot of the proportion of loci rejecting the GTRGAMMA model for the 25 Reid et al. data sets and the 22 phylogenomic data sets. A two-sample t-test shows no significant difference between two sets of data with p-value = 0.96. c) Boxplots of GC content of loci rejecting or accepting the GTRGAMMA model. d) Boxplots of informative sites of loci rejecting or accepting the GTRGAMMA model. e) Logistic regression of whether a locus rejects the GTRGAMMA model (response variable) versus GC content and informative sites (two explanatory variables). The y-axis is the count of data sets with a negative or positive coefficient of GC content (or number of informative sites) estimated in the logistic regression. The bars in dark grey denote the count of data sets for which the coefficient (negative or positive) of GC (or informative sites) is significant (p-value < 0.05) in the logistic regression.

To understand the causes of poor substitution model fit, we investigated the relationship between GC content (and proportion of informative sites) and the rejection of the GTRGAMMA model. A two-sample t-test suggests that the proportions of GC content and informative sites of loci rejecting the GTRGAMMA model are significantly higher (p-value < 0.05) than those for loci that fit the GTRGAMMA model (Figure 2c-d). We fitted a logistic regression for all loci across the 47 data sets, where non-significance (0) or significance (1) of a locus in the intersection test is the binary response variable, and the proportions of GC content and informative sites are two explanatory variables. In the fitted logistic regression, the coefficients of two explanatory variables are significantly negative with p-value < 0.01. We further fit a logistic regression to each of the 47 data sets. The coefficient of GC content is negative/positive for 29/18 data sets, among which 6/1 negative/positive coefficients are significant at the α level of 5% (Figure 2e). Similarly, the coefficient for the number of informative sites is negative/positive for 33/14 data sets, among which 14/0 negative/positive coefficients are significant (Figure 2e). The preponderance of significantly negative coefficients indicates that GC content and the number informative sites are both negatively correlated with the probability of a locus rejecting the GTRGAMMA model, i.e., a higher GC content and/or proportion of informative sites tends to increase the chance of a poor fit of the GTRGAMMA substitution model.

### Testing topologically congruent gene trees

The LRT for TC gene trees rejected the null hypothesis of tree congruence for all 47 empirical data sets with p-value < 0.05. Thus, all empirical data sets in this study strongly favor the alternative hypothesis of incongruent gene trees, a pattern that cannot be adequately explained by gene tree estimation errors. The p-values of the data sets with 10 or more loci are very close to 0, indicating that the assumption of TC gene trees is rarely satisfied for phylogenomic data, which often involve thousands of loci. A two-sample t-test finds no significant difference (p-value = 0.10) in the proportion of loci rejecting the null hypothesis of TC gene trees between the 22 phylogenomic data sets and the 25 Reid et al. data sets (Figure 3a). The topological congruence LRT on individual loci suggests that 38% of gene trees across 47 data sets are statistically incongruent with the concatenation tree (Figure 3b). When the 47 data sets are grouped into 6 categories – mammals (11), birds (11), insects (6), fish (5), reptiles (5), and others (9), an ANOVA analysis finds no significant difference in the proportion of loci rejecting the hypothesis of TC gene trees among 6 groups (Figure 3c). Both the two-sample t-test and ANOVA indicate that the proportion of loci rejecting the hypothesis of gene tree congruence is similar across groups and data sets. A linear regression line was fit for the log scale of the number of incongruent loci rejecting the hypothesis of TC gene trees (*y*) versus the log scale of the number of loci (*x*), i.e., log(y) = 0.87 x log(x) - 0.08 with a significant (p-value < 0.01) positive correlation between log(x) and log(y) (Figure 3d). This result is consistent with the previous conclusion that phylogenomic data sets with more loci are more likely to reject the assumption of TC gene trees; namely, the observed gene tree variation cannot be adequately explained by gene tree estimation error. Moreover, both ANOVA and linear regression analyses suggest a constant and high proportion (38%) of loci rejecting the assumption of TC gene trees across 47 data sets, providing strong evidence for violation of the concatenation assumption of congruent gene trees in phylogenomic data across the tree of life. When the 22 phylogenomic data sets are grouped by data types – CDS (10), EXON (4), UCE (8), the t-tests for pairwise comparisons find no significant difference (p-value = 0.1) for the proportion of loci rejecting the assumption of TC gene trees between the CDS and Exon groups, but the proportions of both groups are significantly (p-value < 0.01) higher than that of the UCE group (Figure 2e). This result indicates that the congruent gene tree assumption of the concatenation model is more likely to hold for the UCE data than for the CDS and Exon data.

**Figure 3:**
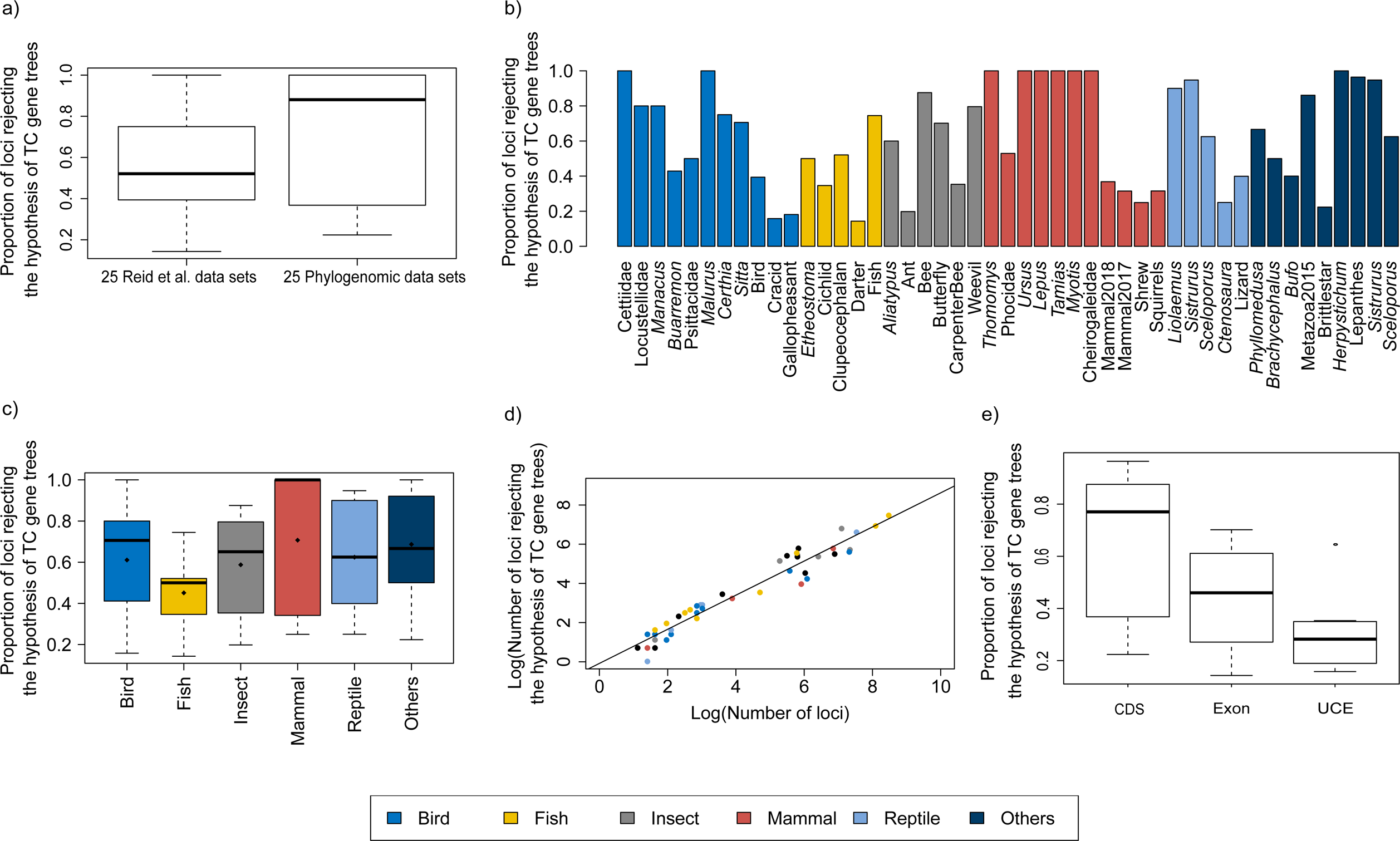
LRT for topologically congruent gene trees. a) A scatter plot of the p-value versus the number of loci rejecting the hypothesis of TC gene trees across 47 data sets. b) Boxplots of the proportion of loci rejecting the hypothesis of TC gene trees for 25 Reid et al. data sets and 22 phylogenomic data sets. A two-sample t-test shows no significant difference (p-value = 0.10) in the proportion of loci rejecting the hypothesis of TC gene trees between the 25 Reid et al. data sets and the 22 phylogenomic data sets. c) Proportion of loci rejecting the hypothesis of TC gene trees across 47 data sets. d) Proportion of loci rejecting the hypothesis of TC gene trees in 6 groups. The 47 data sets fall into 6 groups (bird, mammal, insect, fish, reptile, and others). The filled black diamond represents the average proportion of loci rejecting the hypothesis of TC gene trees in each group. ANOVA finds no significant difference (pvalue = 0.71) in the proportion of loci rejecting the hypothesis of TC gene trees among 6 groups. e) A linear regression line fitted for the log of the number of loci rejecting the hypothesis of TC gene trees (*y*) versus the log of the number of loci (*x*). The regression line is log(y) = 0.87 x log (x) - 0.08, where the coefficient 0.87 is significantly greater than 0 with pvalue < 0.01.

To reduce the potential bias caused by an unfit substitution model, the LRT was only applied to the loci that fit the GTRGAMMA model. We filtered out 27 data sets with < 5 loci that fit the GTRGAMMA model and applied the LRT to the remaining 20 data sets. The null hypothesis of tree congruence was rejected for all 20 data sets with p-value < 0.05. The LRT on individual loci suggests that 46% of gene trees across 20 data sets are statistically incongruent with the concatenation tree, which is lower (but not significantly) than the proportion (48%) when the LRT was applied to all loci of the 20 data sets (Figure A2).

### Bayesian model validation

The coalescent methods have been widely used for estimating species trees from phylogenomic data. Due to computational constraints, however, few studies have evaluated the fit of the MSC to the multilocus sequences. Here, we validate the MSC model using two tests based on Bayesian predictive simulation. The first test (i.e., PPS proposed by Reid et al. (2014)) compares the simulated gene trees with the posterior coalescent gene trees generated with the MSC prior, whereas the second test (i.e., the independent test) compares the posterior coalescent gene trees with the posterior independent gene trees generated with the independent prior. The analysis of 25 Reid et al. data sets found that 8 (32%) data sets failed either or both of two tests (Figure 4a), among which 3 data sets were also found to poorly fit to the MSC by Reid et al (2014; *Certhiidae, Tamias, Aliatypus*). However, the xml input files provided by Reid et al (2014) assumed the HKY and TN93 (+GAMMA) substitution models for all loci. When the *BEAST analyses were rerun with the GTRGAMMA model, only two data sets provided were rejected by either or both of two tests (Table 2). Thus, the choice of substitution models has major effects on the fit of the MSC. In addition, Bayesian model validation for the 14 phylogenomic data sets for which the GTRGAMMA model was a good fit found that 12 data sets failed the first test and all 14 data sets failed the second test.

**Figure 4:**
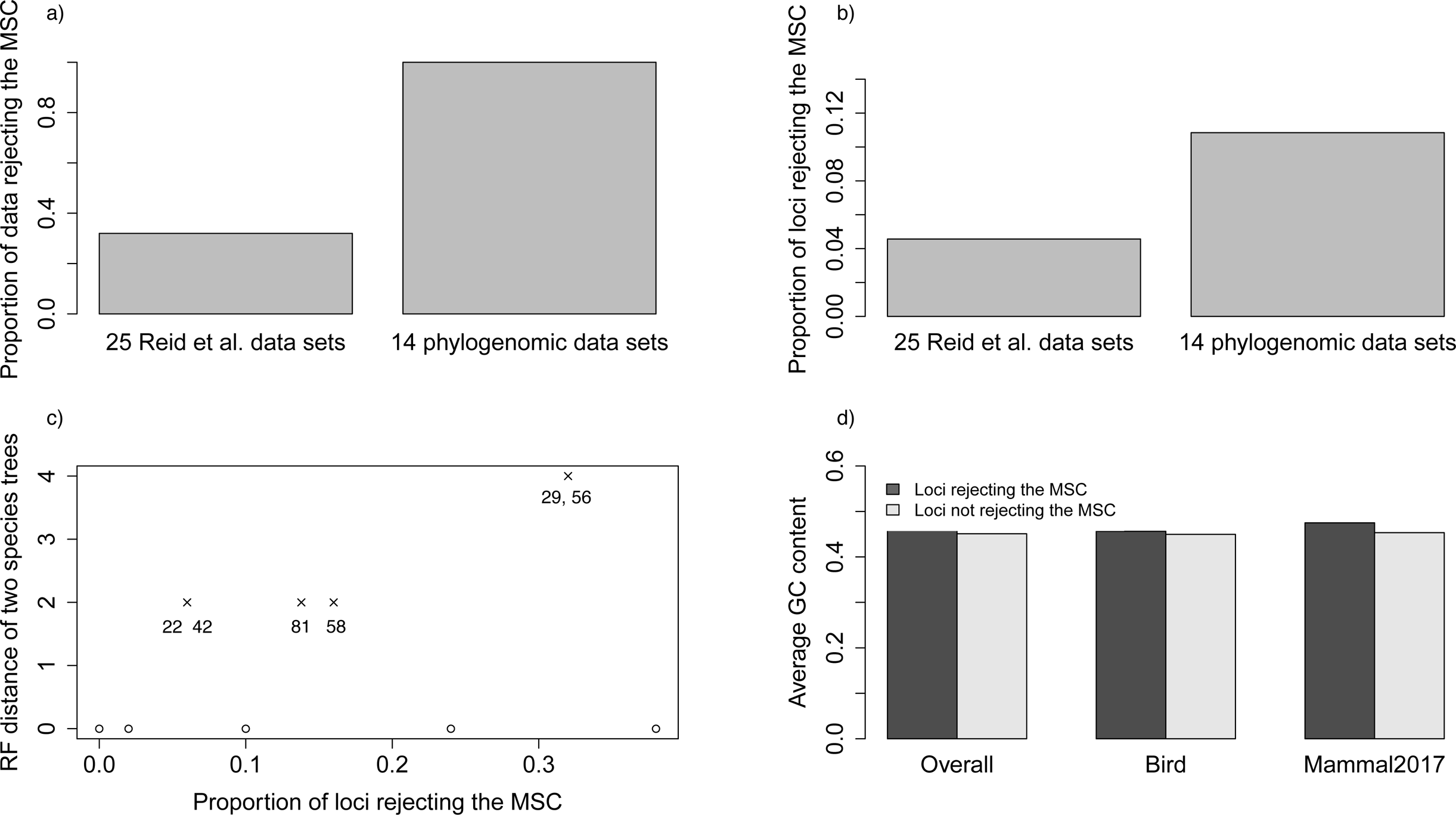
Bayesian model validation. The MSC model is validated by two tests. The first test proposed in Reid et al. (2014) compares the simulated and posterior coalescent gene tree. The second test compares the gene trees estimated with the coalescent prior and those estimated with independent prior. The MSC model is a poor fit to the data if either or both of the two tests are rejected. a) The proportion of data sets rejecting the MSC model. 32% of 25 Reid et al. data sets and 100% of 14 phylogenomic data sets reject the MSC model. b) The tests were applied to each locus to find the proportion of loci rejecting the MSC model. The proportion (10.88%) of loci rejecting the MSC for 22 phylogenomic data sets is significantly (p-value < 0.01) higher than the proportion (4.17%) of loci in 25 Reid et al. data sets that reject the MSC model. c) A scatter plot of the RF distance versus the proportion of loci rejecting the MSC model. The RF distance between two species trees built from all loci and only loci that fit to the MSC is calculated for 14 phylogenomic data sets; RF = 0 for 9 data sets, RF = 2 for 4, RF = 4 for 1 data set. The number below each point is the bootstrap support value on the corresponding incongruent branch in two species trees. Note that two points with RF = 2 are overlapping, and two values below the point with RF = 4 (i.e., two incongruent branches) are the bootstrap support values on two incongruent branches. d) Testing the association between GC content and poor fit of the MSC. We calculated the average GC content of loci rejecting or accepting the MSC for two phylogenomic data sets with large numbers of loci rejecting the MSC (Birds and mammal2017). We performed a two-sample t-test for the overall average GC content (combining two data sets) and the average GC content of each of two data sets, respectively. All three tests found no significant difference (p-value > 0.3) in GC content between loci rejecting the MSC and loci accepting the MSC.

**TABLE 2:**
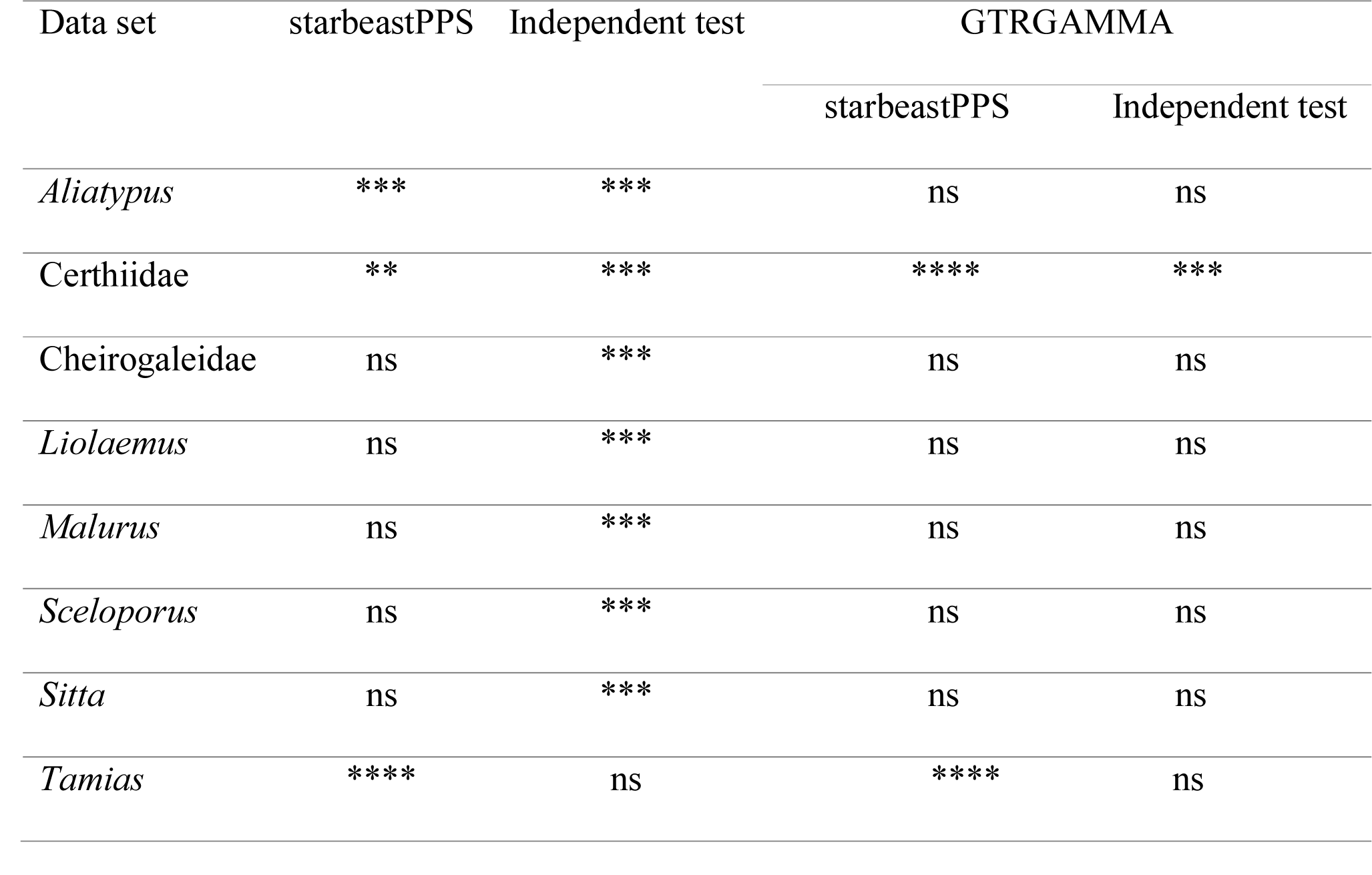
Bayesian model validation for the 25 data sets in Reid et al. (2014). The validation analysis involves two tests - starbeastPPS and the independent test. Significance symbols *<0.05, **<0.025, ***<0.01, ****<0.001, and ns denotes nonsignificant. The last column GTRGAMMA indicates that the Bayesian model validation analyses assuming HKY and TN93 + GAMMA were reconducted with the GTRGAMMA model.

Thus, the MSC model failed to fit all 14 phylogenomic data sets (Figure 4a). Since these 14 data sets have more loci than the Reid et al. data sets, it implies that phylogenomic data with more loci are more likely to reject the MSC. In addition, Bayesian model validation for individual loci of the 14 phylogenomic data sets found that 10.88% of loci rejected the MSC (Figure 4b), significantly higher (p-value < 0.01) than the proportion (4.17%) for the 25 Reid et al. data sets, indicating that the probability of a locus rejecting the MSC increases as the number of loci grows.

To evaluate the effect of loci rejecting the MSC on species tree estimation, two species trees were reconstructed from all loci and only loci that fit to the MSC for each of 14 phylogenomic data sets. A majority (9) of 14 data sets produced two identical species trees (i.e., RF = 0), whereas RF = 2 for 4 data sets (Ant, Cracid, Mammal2017, Squirrel) and RF = 4 for 1 data set (Figure 4c). Note that RF = 2 or 4 indicates only 1 or 2 conflicting branches in two species trees. Since the incongruent branches are not strongly supported (bootstrap support values < 60), the conflict between two different species trees is not significant. This analysis suggests that including loci that fail to fit the MSC has little impact on species tree estimation, when a small portion (10.88%) of loci rejects the MSC.

To investigate the association between GC content and poor fit of the MSC, we calculated the average GC content of loci rejecting or accepting the MSC for two phylogenomic data sets for which the number of loci rejecting the MSC is large (Birds and mammal2017, in which the number of loci rejecting the MSC is 10 and 17, respectively; Figure 4d). Other phylogenomic data sets contain insufficient number of loci rejecting the MSC for the analysis. A two-sample t-test for the overall average GC content (combining two data sets) and the average GC content of each of two data sets unanimously found that the difference in GC content between loci rejecting the MSC and loci accepting the MSC was not significant (Figure 4d), showing little evidence for a positive association between high GC content and poor fit of the MSC.

### Bayesian model comparison for MSC versus concatenation

Bayesian model comparison was applied to 31 data sets for which there was no missing data, including 17 Reid et al. data sets and 14 phylogenomic data sets. The BFs (on logarithmic scale) of 22 data sets are greater than 100 and the BFs of the remaining 9 data sets are between 10 and 100. Overall, the high BFs imply that all 31 data sets strongly favor the MSC rather than the concatenation model (Figure 5). This Bayesian model comparison analysis is consistent with the LRT results for congruent gene trees, which reject the concatenation assumption of congruent gene trees and thus favor the MSC for all 47 data sets.

**Figure 5:**
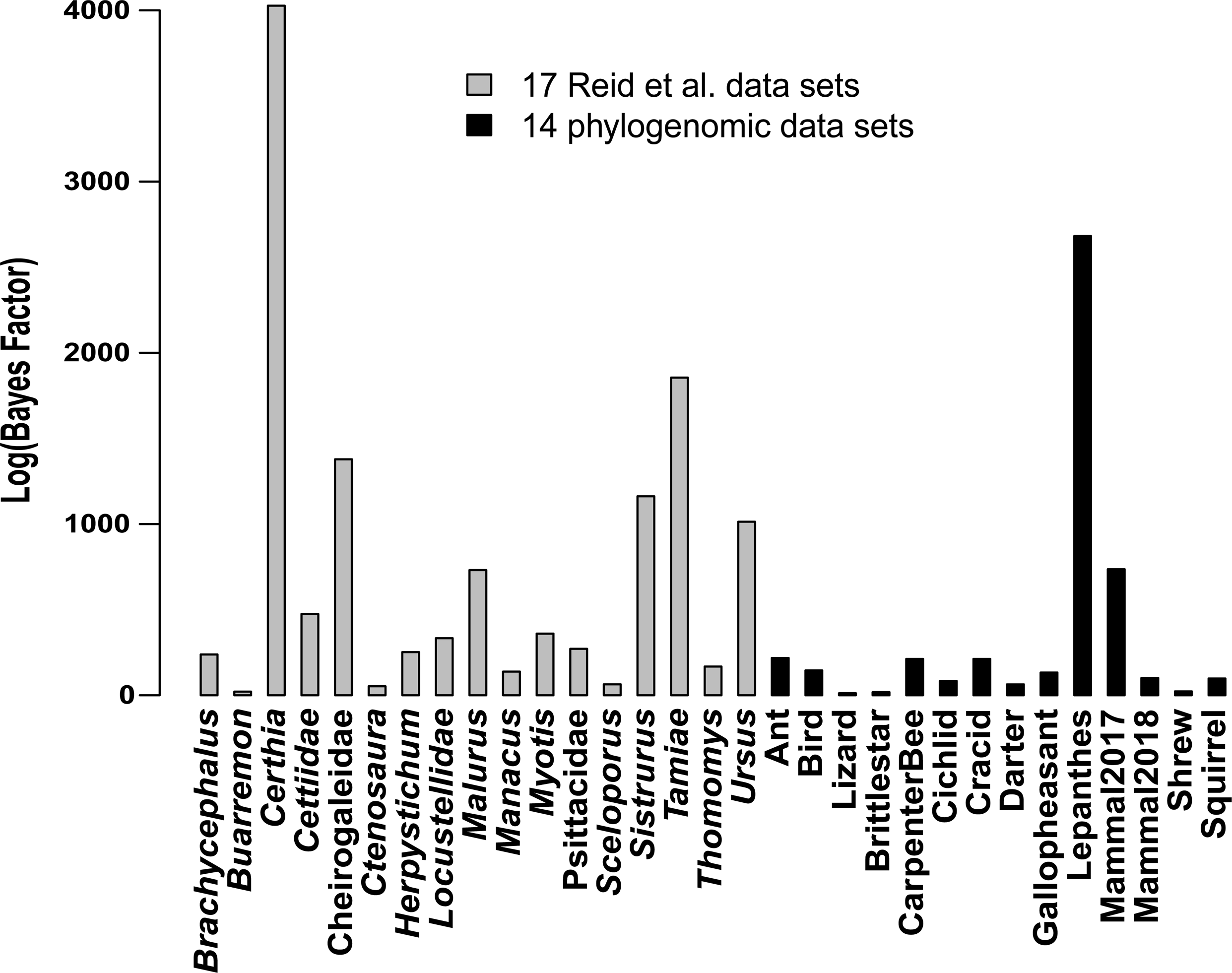
Bayesian model comparison. The log-scale Bayes factors of coalescent vs concatenation (unlinking substitution model parameters) for 31 data sets, including 17 Reid et al. data sets and 14 phylogenomic data sets.

To demonstrate the impact of model assumptions on species tree inference, species trees were estimated for 4 phylogenomic data sets (Cracids 23 species and 430 loci, Gallophesants 18 species and 1479 loci, Lizard 29 species and 1852 loci, Shrews 19 species and 1112 loci, Table 1) using concatenation and a coalescent method NJst. We found 3 data sets (Cracids, Gallophesants, Lizard) for which the concatenation trees were significantly incongruent with the corresponding NJst trees (Figure A3), indicating that different models may yield conflict species trees. In the subsampling analysis, the average proportion of significantly incongruent concatenation trees across 4 data sets is 0.19, much higher than the average proportion (0.008) of significantly incongruent NJst trees (Figure A4), suggesting that concatenation is more likely than coalescent methods in producing wrong relationships with high bootstrap support values.

## DISCUSSION AND CONCLUSIONS

Model validation and comparison are essential to accurate phylogenetic inference for genome-scale sequence data. Many recent disputes about the utility of the MSC model in phylogenomics have rested on perceived model violations of the MSC, rather than direct tests of the explanatory power of the MSC versus concatenation. Here we developed and implemented a set of statistical tests to evaluate the adequacy of substitution models, the concatenation model, and the MSC model. In particular, we tried to distinguish two possible sources of rejection of the MSC in empirical data sets: rejection due to violation of the substitution model and rejection due to violation of the MSC. The LRT results for congruent gene trees reveals strong evidence in 47 data sets against the concatenation assumption of congruent gene trees across loci. Crucially, this test suggests that the gene tree variation is real and cannot be explained simply by gene tree estimation error, a point of increasing concern among skeptics of the MSC model (Richards *et al*. 2018). This result is consistent with the subsequent Bayesian model comparison analysis, which unanimously favored the MSC over concatenation for all phylogenomic data sets under consideration. Moreover, the proportion of gene trees significantly deviating from the concatenation tree (38%) is consistently high across taxonomic groups (bird, fish, mammal, insect, reptile, and others), and our linear regressions suggest that the concatenation assumption of congruent gene trees is more seriously violated as the number of loci continues to grow in phylogenomic data across the tree of life (Bravo *et al*. 2019).

The fact that Bayesian model comparison strongly favors the coalescent over concatenation does not necessarily validate the use of the MSC model for analyzing phylogenomic data. In our Bayesian model validation, the MSC model is a good fit for the majority data sets in Reid et al. (Reid *et al*. 2014), but the choice of substitution models has a strong influence on the fit of the MSC model. To alleviate the effect of substitution models, we applied Bayesian model validation to the loci of 14 phylogenomic data sets that fit the assumed substitution model. The MSC failed to fit all 14 phylogenomic data sets, but the proportion of loci rejecting the MSC was only 11%, significantly smaller than those for substitution models (44%) and concatenation models (38%). Thus, deficiencies in the fit of data to substitution models and to concatenation models appear to be much more severe than fit to the MSC model, suggesting that more attention should be given to appropriately modeling the evolution of single nucleotides (i.e., substitution models) and gene tree variation, though continuous efforts for improving models at the level of both sequences and gene trees are ultimately desirable.

An empirical study of placental mammals (Romiguier *et al*. 2013) suggested that GC-rich regions are associated with high recombination rates which may be problematic for species tree inference under the MSC model. Our analysis, however, finds no convincing evidence for the positive association between high GC content and poor fit of the MSC. Instead, we find that high GC content is strongly associated with poor fit of substitution models. Thus, the shifting phylogenetic relationships of placental mammals for GC-rich regions found by Romiguier et al. (2013) may be caused not by poor fit of the MSC, but the conflicts among gene trees due to poorly fitted substitution models.

Stochastic models for phylogenomic data should consider the cumulative effect of the mutation process of nucleotides (molecular evolution) and biological processes rooted in population genetics that have played important roles in the evolution of species. Some have argued (Edwards 2009; Liu et al. 2015) that, among the relevant biological processes, the coalescence process, which assumes random drift, should serve as the null model, and other biological factors, such as gene flow and hybridization, can be added to the null model if the null model cannot adequately explain the observed gene tree variation. As the number of loci continues to increase, some loci are bound to reject the MSC model. Brown and Thomson (2017) suggests that a small number of extremely influential loci can significantly change the estimates of phylogenetic trees. Our analysis indicates that if the loci rejecting the MSC (or concatenation) only account for a small proportion of the empirical data (e.g. 11% in this study), the MSC model (or concatenation) can still be applied to entire data sets or to data sets purged of the loci that violate the MSC. On the other hand, a large number of loci rejecting the MSC (or concatenation) suggests that additional biological phenomena may have occurred and must be added to the stochastic model when analyzing such data sets.

Mathematical models are variably robust to assumption violations. Several authors have identified numerous putative biological violations of the MSC in empirical data sets, including recombination within loci, pseudo-concatenation of loci such as occurs in transcriptome data as well as natural selection (Gatesy and Springer 2014; Scornavacca and Galtier 2017). However, even in these data sets, despite numerous putative violations of the MSC, the MSC is a better fit than concatenation, suggesting that violations of the MSC may not recommend falling back on concatenation as an alternative method of analysis (Liu *et al*. 2015; Edwards *et al*. 2016). The analysis of empirical data in this study suggests that, although 11% of loci reject the MSC, there is no significant difference between the species trees estimated from all loci and only loci that fit the MSC model. Thus, gene-tree-based coalescent methods are robust to a certain degree of violation of coalescent assumptions. In this study, model validation and species tree analyses were conducted on reduced phylogenomic data sets of 50 loci and 10 species each. Our analyses suggest that the concatenation assumption of congruent gene trees rarely holds for phylogenomic data with more than 10 loci. Thus, for large phylogenomic data sets, model comparison analyses are expected to consistently and more strongly favor the coalescent model over the concatenation model. As the number of loci increases, model validation analysis is more likely to reject the MSC. Adding species above the level of genus in phylogenomic data, however, will introduce additional gene tree variation caused by gene flow, gene duplication/loss, etc. Thus, unlike increasing numbers of loci, growth in the number of taxa in phylogenomic data is likely to increase the proportion of loci rejecting the MSC. When the majority of loci do not support the MSC, the coalescent methods will eventually fail to accurately reconstruct species trees from the full phylogenomic data, even though model comparison is still in favor of the MSC over the concatenation model. In such cases, MSC network models, in which gene flow and lineage merging are incorporated, may better fit phylogenomic data sets than the standard MSC model (Wen *et al*. 2016; Bastide *et al*. 2018).

## SUPPLEMENTARY MATERIAL

The alignments of the input data sets for the substitution model, TC gene trees, model validation, and model comparison analyses are available from the Dryad Digital Repository: http://dx.doi.org/10.5061/dryad.7q6q3s0

## ACKNOWLEDGEMENTS

We thank Noah Reid for sharing 25 multilocus sequence data and the xml *BEAST input files and Bryan Carstens for the valuable comments. We thank Georgia Advanced Computing Research Center (GACRC) for their computing resource.

